# The impacts of hydropower on freshwater macroinvertebrate richness: A global meta-analysis

**DOI:** 10.1101/2021.02.05.429909

**Authors:** Gabrielle Trottier, Katrine Turgeon, Daniel Boisclair, Cécile Bulle, Manuele Margni

**Affiliations:** CIRAIG, Département de mathématique et génie industriel, Polytechnique Montréal, Montréal, Québec, Canada; ISFORT, Université du Québec en Outaouais, Ripon, Québec, Canada; Département de sciences biologiques, Université de Montréal, Montréal, Québec, Canada; CIRAIG, Département de stratégie, responsabilité sociale et environnementale, École des sciences de la gestion, Université du Québec à Montréal, Montréal, Québec, Canada; Institute of Sustainable Energy, HES-SO Valais-Wallis, Sion, Switzerland

## Abstract

Hydroelectric dams and their reservoirs have been suggested to affect freshwater biodiversity. However, studies investigating the consequences of hydroelectric dams and reservoirs on macroinvertebrate richness have reached opposite conclusions. We carried out a meta-analysis devised to elucidate the effects of hydropower dams and their reservoirs on macroinvertebrates richness while accounting for the potential role played by moderators such as biomes, impact types, study designs, sampling seasons and gears. We used a random and mixed effect model, combined with robust variance estimation, to conduct the meta-analysis on 72 pairs of observations (*i*.*e*., impacted versus reference) extracted from 17 studies (more than one observation per study). We observed a large range of effect sizes, from very negative to very positive impacts of hydropower. However, according to this meta-analysis, hydropower dams and their reservoirs did not have an overall clear, directional and statistically significant effect on macroinvertebrate richness. We tried to account for the large variability in effect sizes using moderators, but none of the moderators included in the meta-analysis had statistically significant effect. This suggests that some other moderators, which were unavailable for the 17 studies included in this meta-analysis, might be important (*e*.*g*., temperature, granulometry, wave disturbance and macrophytes) and that macroinvertebrate richness may be driven by local, smaller scale processes. As new studies become available, it would be interesting to keep enriching this meta-analysis, as well as collecting local habitat variables, to see if we could finally draw statistically significant conclusions about the impacts of hydropower on macroinvertebrate richness.

## Introduction

Freshwater ecosystems are vital resources for humans and support a biota that is rich, sensitive and characterized by a high level of endemicity [1]. Ecosystems functions and integrity often depend on biodiversity, which can be described by three indices: species richness (*i*.*e*., number of species), community assemblage (*i*.*e*., proportions of different species or taxonomic groups in the community) and functional diversity (*i*.*e*., variability in organisms’ traits that can influence ecosystem functioning; [1]). For millennia, humans have used freshwater ecosystems, through water extraction for drinking and irrigation purposes, water regulation for hydropower production, flood control and recreation [1] but these usages often come with a cost on freshwater ecosystems biodiversity [2,3].

Hydroelectric dams and the creation of reservoirs, at all stages of their life cycle (*i*.*e*., from the construction, to operation and decommission of a dam), can affect freshwater biodiversity [4]. Dams create a physical barrier, which can impair the natural flow of water, sediments and nutrients [5,6] and limit the movement of organisms [7]. The alteration of the natural hydrological regime can affect freshwater biodiversity through various biological mechanisms (*i*.*e*., mortality through desiccation, mismatch timing in life history strategies, lotic to lentic community changes, reduction/extirpation of endemic and specialist species; [8,9]) and through degraded water quality (anoxic or hypoxic releases [dissolved oxygen], hypolimnetic or epilimnetic releases [temperature], pH, organic carbon, turbidity; [4,10–13]).

Studies investigating the impact of hydropower on macroinvertebrate richness drew contrasting conclusions. Some studies reported that macroinvertebrate richness is negatively impacted by hydropower, through general flow regulation [14–17] and water level fluctuation (or drawdown; [18–23]). Others observed higher macroinvertebrate richness downstream of a dam [24,25] or in regulated rivers (as opposed to natural ones; [26]). Finally, Marchetti et al. [27] found little difference between (*i*.*e*., dam-induced permanent low flow) and “natural-like” flows (*i*.*e*., high flows in winter and spring, low flows in summer and falls). A meta-analysis could allow to elucidate the patterns and interaction that may exist between hydropower, macroinvertebrate richness, and the context in which studies have been conducted.

Many challenges can be encountered when conducting a meta-analysis, and many ecological facets of studied ecosystems can influence the magnitude and significance of human impacts [28], along with study-specific methodological characteristics (*e*.*g*., different study design). The influence and the variability brought about by these facets and characteristics can be accounted for through variables, also called moderators in meta-analysis [29]. For instance, the location of each study site can influence the observed effects [28,30]. A latitudinal biodiversity gradient is a good example of the influence of spatial location. Species richness is known to be highest in the tropics and lowest at the poles [31,32]. Hydropower can lead to different types of impacts. A study can analyze the impacts of hydropower upstream of a dam, that is in the reservoir, or downstream of a dam. Impacts also varies depending on the type of water management in place, storage reservoir with winter water level drawdown, hydropeaking or typical run-of-river hydrological regime. These variations in studies, along with the location under study, can introduce variability and heterogeneity in the results, which can be accounted for through moderators. The experimental design can also influence how human-induced impacts magnitude are reflected in a study [28]. As demonstrated in Christie et al. [33], different sampling designs may affect the conclusion of a study. Using simulations, they demonstrated that Before-After (BA), Control-Impact (CI; analogous to space-for-time substitution) and After designs are far less accurate than Randomized Controlled Trials (RCT) and Before-After Control-Impact (BACI) designs. RCT and BACI are much harder to implement in ecology because true randomization can be difficult with larger scale designs and getting data before the impacts or human intervention is sometimes impossible. Thus, we must account for the effect of the experimental design on a study outcome, especially in a meta-analysis, where the effects of multiple different studies are combined. Sampling season can also influence the results across studies, as macroinvertebrate communities differs in terms of abundance and diversity throughout the year (*i*.*e*., maximum diversity in late summer and autumn versus underrepresented diversity in spring and early/mid-summer; [34]). At a more local scale, the habitat stratum that is sampled is also most likely to influence the results [28], especially when studying macroinvertebrates. These organisms possess characteristics that make them highly adapted to their environment [35] and because lakes, reservoirs and river beds are so heterogeneous, macroinvertebrates are often patchily distributed, requiring extensive sampling [36]. Thus, the type of sampling gear used to sample will likely affect the type of organisms inventoried in each study.

The objective of this manuscript is to conduct a meta-analysis about the impacts of hydropower dams and their reservoirs on macroinvertebrates richness while accounting for a series of moderators defining the context of the studies included. The moderators included in this manuscript are the following: 1) biomes (*i*.*e*., boreal, temperate, and tropical), a proxy for location/latitudinal gradient, 2) type of impact, which is reflected by the position of a sample in relation to the dam (*i*.*e*., upstream or downstream of the dam). Downstream stations are impacted by a reduced flow and hydropeaking dynamics, whereas upstream stations are impacted by drawdown and water level fluctuations due to reservoir management, 3) type of study designs such as cross-sectional (*i*.*e*., reference natural lake versus impacted reservoir) and longitudinal spatial gradient (*i*.*e*., upstream of a dam [reservoir] versus downstream of a dam [river]), which are two different variants of CI study design, 4) sampling seasons (*i*.*e*., spring, summer, fall, winter) and 5) sampling gears, a proxy for habitat stratum (*i*.*e*., grabs and nets).

## Methodology

A meta-analytic framework was used to assess the impacts of hydropower on macroinvertebrate richness, for different biomes, impact types, study designs, seasons and sampling gear. A meta-analysis is a method of research synthesis that is based on expressing results from multiple studies that share a given research subject (but different sampling strategies, sample sizes or sampling gears) on a common comparison scale, and aims to integrate findings, establish generalizations or resolve conflicts [37]. Whereas a research synthesis is usually performed qualitatively, a meta-analysis is a powerful, informative and unbiased tool that allows the quantification of generalizations through various statistical methods [37]. It statistically combines the magnitude and the direction of results or outcomes (later referred to as effect sizes) of multiple studies sharing a common research objective [37]. For example, when comparing richness between an impacted and a natural habitat, the magnitude of the difference in richness (*i*.*e*., delta) and its direction (*i*.*e*., either gain or loss of taxa) constitute the effect size. Meta-analysis has many advantages over other methods (*i*.*e*., narrative review, vote counting). Namely, it takes into account sample size and statistical power from each study (with potentially different sampling strategies), assesses the magnitude and statistical significance of the mean effect size, and allows the analysis of multiple sources of variation among studies (*i*.*e*., moderators; [37]). For those reasons, we chose this method over the others.

Achieving our objective requires to first establish a research strategy, second to do a data collection comprising information regarding macroinvertebrate richness in hydropower impacted habitat versus reference ones, biome, the type of impact, study design, sampling seasons and gear. Third, it requires to compute effect sizes for each study and finally, combine them to assess if the mean effect size is significantly different from zero [37] and if the presence of other moderators can influence these results.

### Research strategy

In this study, we used the PRISMA (Preferred Reporting Items for Systematic and Meta-Analyses) methodology, flow diagram (S1 Fig) and checklist (S1 Table) proposed by Moher et al. [38], to report systematic literature reviews and meta-analyses. A systematic literature review was conducted using the Web of Science Core Collection database, which includes all journals indexed in Science Citation Index Expanded (SCI-EXPANDED), Social Sciences Citation Index (SSCI), Arts & Humanities Citation Index (A&HCI), Conference Proceedings Citation Index – Science (CPCI-S), Conference Proceedings Citation Index – Social Science & Humanities (CPCI-SSH) and Emerging Sources Citation Index (ESCI; [39]). The research strategy was constrained between 1989 and 2019 and contained a combination of the four following field of research: 1) hydropower (hydropower OR hydroelectric* OR dam OR dams OR reservoir* OR impound* OR run-of-river OR “run of river” OR drawdown* OR hydropeak* OR dam* OR “water level fluctuation” OR “water-level fluctuation” OR “water level variation” OR “water-level variation” OR “water level regulation” OR “water-level regulation” OR “water level manipulation” OR “water-level manipulation” OR “water management”), 2) biodiversity (“biodiversity” OR “richness”), 3) freshwater ecosystems (“freshwater” OR “aquatic”) and 4) aquatic insects (“*invertebrate*” OR “*benth*” OR “insect*” OR “arthropod*”). Research strategy also excluded all studies pertaining to beaver and agricultural dams (NOT “beaver*” NOT “agricult*”). The research strategy resulted in 408 research articles, as per December 16^th^ 2019.

The results were sorted by Web of Science relevance, which ranks records based on the number of search terms found in each record [40], and extracted as a list to further evaluate the relevance of every studies based on a list of criteria. The following criteria were applied to assess the inclusion of any study in the meta-analysis: 1) the study had to refer specifically to hydropower related impacts (*i*.*e*., reservoir, run-of-the-river or hydropeaking, multi-purpose reservoirs were also checked for hydropower impacts), 2) the scope of the study had to address freshwater ecosystems and macroinvertebrates and 3) the study had to be empirical (*i*.*e*., excluding literature reviews, modelling exercises) and provide an explicit richness, error and sample size value for both a reference and impacted site (*i*.*e*., cross-sectional studies [reference versus impacted]) or gradient of impact (longitudinal spatial gradient studies [upstream of the dam/reservoir versus one or multiple sites downstream of the dam]). Out of the 408 research studies that resulted from the research strategy, only 17 met the above criteria (see geographical disposition of studies in Fig 1).

**Fig 1.**
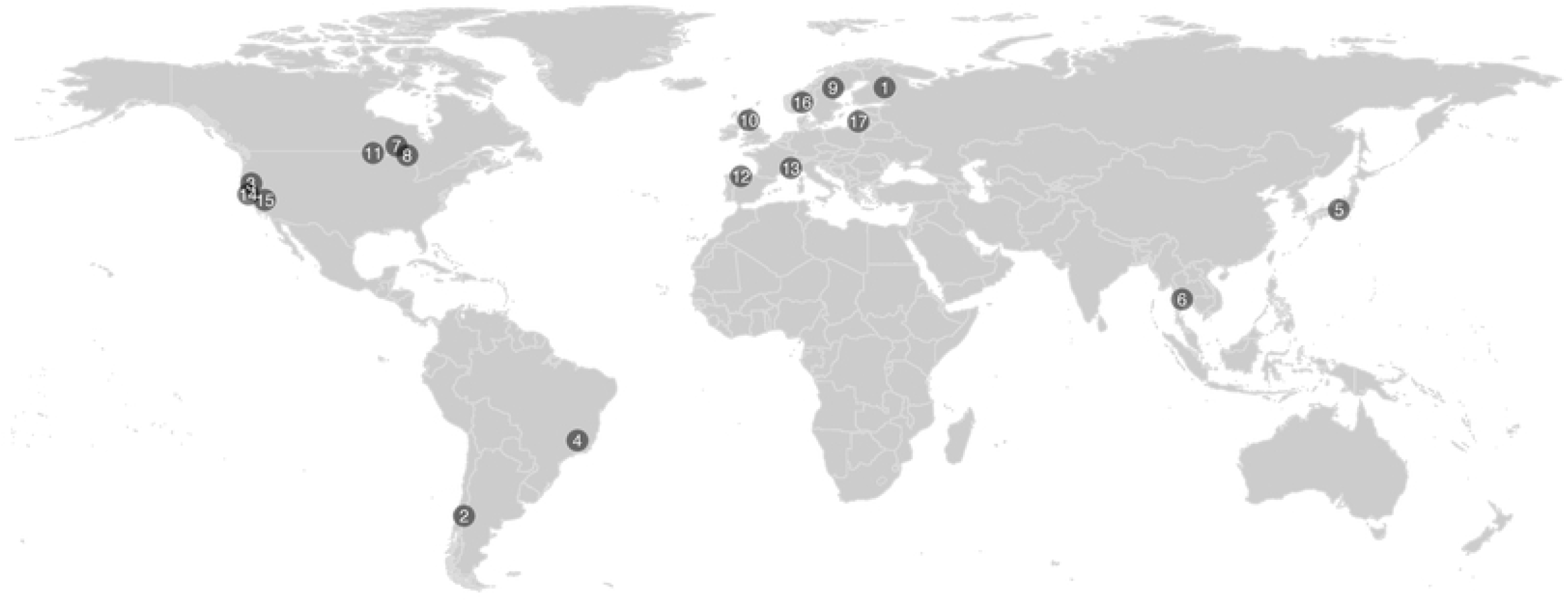
World map showing the geographical disposition of the studies used in this meta-analysis; [1] Aroviita and Hämäläinen (2008), [2] Valdovinos et al. (2007), [3] Marchett i et al. (2011), [4] Molozzi et al. (2013), [5] Takao et al. (2008), [6] Kullasoot et al. (2017), [7] White et al. (2011), [8] Smokorowski et al. (2011), [9] Englund and Malmqvist (1996), [10] Jackson et al. (2007), [11] Kraft (1988), [12] Mellado-Dìaz et al. (2019), [13] Bruno et al. (2019), [14] Milner et al. (2019), [15] Steel et al. (2018), [16] Schneider and Petrin (2017) and [17] Vaikasas et al. (2013).

### Data collection

Richness metrics and moderators were collected for each of these 17 studies (S2 and S3 Tables). We extracted richness (*i*.*e*., number of taxa), error measure (*e*.*g*., standard deviation), sample size (*i*.*e*., number of reference observations and impacted observations used to compute richness and its associated error) and a suite of moderators such as biome (*i*.*e*., boreal, temperate and tropical; [41]), type of impact (*i*.*e*., water level fluctuations upstream, due to reservoir management, or flow regulation downstream due to dam operations), type of study, (*i*.*e*., cross-sectional [reference natural lake versus impacted reservoir] or longitudinal spatial gradient [reference; upstream of the dam/reservoir versus impacted; downstream of the dam]), sampling season (*i*.*e*., spring [March to May], summer [June to August], fall [September to November] and winter [December to February], according to the hemisphere) and sampling gear (*i*.*e*., net, grab or colonization basket). These moderators were chosen based on the availability of said moderators in each of the 17 studies, their potential influence on macroinvertebrate richness and complemented with expert judgement. In studies where meaningful data were presented exclusively in graphical format, values were extracted using Engauge Digitizer 10.4 [42].

### Effect size

To compute effect sizes, we calculated the standardized mean differences, also called Cohen’s *d* – which expresses the distance between two means (*i*.*e*., impact and reference) in terms of their common standard deviation [43]. For most studies – except Takao et al. [15] and Schneider and Petrin [44], we computed at least two effect sizes per study, leading to a total of 72 effect sizes, with a certain level of within-study dependency (*i*.*e*., effect sizes in one study are not entirely independent from each other). Cohen’s *d* is calculated as per equation 1 [43]:

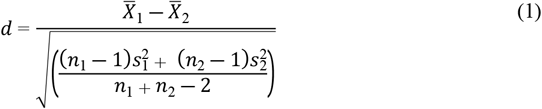

where 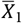 and 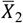 are the mean richness, 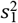and 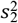 are the standard deviation (SD) and, *n*_1_ and *n*_2_ are the number of observations used to compute the mean and SD for impacted and reference samples, respectively. The common variance (*V*_*d*_) associated with the effect size (*d)* is calculated using equation 2:

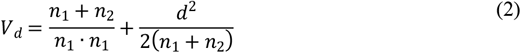

In the case of small sample size (< 20 studies), a correction factor is applied to Cohen’s *d* to reduce the positive bias (negligible with bigger sample size) and to provide a better estimate. The corrected effect size is then called a Hedges’ *g* (Hedges, 1981). A small sample correction factor (*J*) was computed using equation 3:

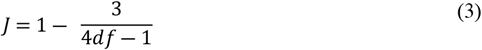

where *df* refers to the degrees of freedom (*n*_1_ + *n*_2_ ―2). Thus, the corrected effect size *g* and variance *V*_*g*_ are calculated following equations 4 and 5, respectively:

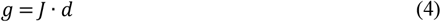

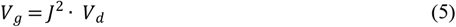

A positive *g* means the impacted environment or sample has more richness in comparison to a reference environment or sample. The *metafor* package [29] was used to compute the effect sizes and sampling variances (*i*.*e*., *escalc* function) and the *ggplot2* package [46] was used to graphically visualize the results of the meta-analysis.

## Data analysis

### Publication bias

Studies with large significant results are more likely to be published than studies with non-significant results, this is called publication bias [37]. A funnel plot was used to evaluate the presence of a publication bias in the meta-analysis [47], and a regression test for funnel plot was used to detect potential asymmetry (*i*.*e*., if only large significant studies, range of outcomes is not well represented and studies with non-significant results might not even be present; [48]). The *metafor* package [29] was used for asymmetry analysis (*i*.*e*., *funnel* and *regtest* functions).

### Heterogeneity

Specific study characteristics (*e*.*g*., biome) and dissimilarities in methodologies among studies (*e*.*g*., study design) can introduce variability, or heterogeneity (not only due to within study sampling error), among true effect sizes [29,49]. It is interesting to examine this heterogeneity and identify the different moderators and their relative contributions to the magnitude and direction of these effect sizes [37]. We evaluated the statistical significance and the magnitude of the heterogeneity of effect sizes among studies (*i*.*e*., heterogeneity analysis) using, respectively, the *Q* statistic and *I*^*2*^ index [49]. The *metafor* package [29] was used for heterogeneity analysis (*i*.*e*., *rma* function).

### Random and mixed-effect model: dealing with heterogeneity

Two types of meta-analytic models can be used in a meta-analysis, fixed or random. A fixed model considers only the studies included in the meta-analysis and within study sampling variability, not between studies [37]. No inference can be made outside this set of studies (*i*.*e*., conditional inferences; [29]). A random-effect model considers the set of analyzed studies as a sample of a larger population of studies [29]. Thus, it allows the researcher to make inferences regarding what would be found if an entire new meta-analysis, with a different set of studies, was performed (*i*.*e*., unconditional inferences; [29]). It also allows two sources of variation, within and among studies [37]. Such an approach is especially appropriate when dealing with heterogeneity among studies [49] and with a mixed-effect model approach, it is also possible to include moderators, which can account for some of that heterogeneity [29]. Here, a random and mixed effect modelling is preferred to a fixed modelling approach since 1) a significant amount of heterogeneity was found in the previous heterogeneity analysis and 2) because it offers the possibility to model and explain some of that heterogeneity using moderators [43].

### Robust variance estimation: dealing with dependency

If our effect sizes were all independent from each other, we could have simply used a random and mixed effects model. However, because this meta-analysis is dealing with multiple effect sizes per study, where observations are not methodologically and spatially independent from each other, it is inappropriate to use a regular meta-analysis approach (*i*.*e*., random and mixed effects model), where the effect sizes are assumed to be independent. One way to account for dependency of effect sizes is to combine the random and mixed effect model with the robust variance estimation (RVE) method. The RVE estimates the overall effect size over studies using a weighted mean of the observed effect sizes [50]. It doesn’t require knowledge about the within-study covariance, it can be applied to any type of dependency and effect sizes, it simultaneously accommodates for multiple sources of dependencies, it does not require the effect sizes to comply to any particular distribution assumptions, it leads to unbiased fixed effects and standard errors estimates and can also give an estimate of among-study variance [50,51]. Because the most common source of dependence within the effect sizes in this meta-analysis is the correlated nature of the observations (*i*.*e*., multiple measures within a study; methodological and spatial correlation) and not the hierarchical nature (*i*.*e*., common nesting structure between studies; a sample within a transect, within site and within a lake), a correlated effects weighting method was used [51]. Thus, we used a RVE based on a correlated effects model and adjusted for small sample size (< 40 studies; [51]). Finally, a sensitivity analysis was computed to assess the effect of a varying rho (*ρ*) value, which is a user-specified value of the within-study effect sizes correlation (*i*.*e*., the correlation between two samples taken in the same water body in one specific study – spatial and methodological correlation; [51]). The *robumeta* package [51] was used to fit the RVE meta-regression model (*i*.*e*., *robu* function) and to compute the sensitivity analysis (*i*.*e*., *sensitivity* function). All statistical analyses were made using R version 3.0.2 [52].

## Results

### Methodological results

The purpose of this first set of results is to validate our methodological approach and choices, they will not be the subject of discussion. No statistical asymmetry was observed in the funnel plot (z = −1.0707; P = 0.2843; S2 Fig), a wide range of results and significance levels were represented by the studies included in this meta-analysis. The total heterogeneity among the effect size was statistically significant (*Q*_df_ = 557.67; P < 0.0001), which indicated greater total heterogeneity than expected by the sampling error alone. The estimated amount of this heterogeneity among the effect sizes was *T*^2^ = 3.78; 95% confidence interval [CI] = 2.75-6.15. Of that total variability, a large amount (*I*^2^ = 90.13%; CI = 86.91-93.69) was due to true heterogeneity between the studies, rather than just methodological variability. Thus, further examination of the heterogeneity is warranted and will be done through the analysis of multiple moderators.

### Meta-analysis results

The meta-analysis of 17 studies (72 pairs of observations; S1 Table) suggests that hydropower dams and reservoirs did not have a statistically significant effect on macroinvertebrate richness. The mean effect size (*i*.*e*., Hedge’s *g*) estimate of our RVE model was −0.864 (95% CI = −1.87 to 0.144; p = 0.088), without accounting for the different moderators (Fig 2). The large confidence interval overlapping zero indicates that the mean effect size is not statistically significant and highlights a wide range of effect sizes across studies. The sensitivity analysis shows that the estimates of the mean effect size and standard errors, as well as the estimate of between-study variance in study-average effect sizes (*τ*^2^), are relatively insensitive to different value of *ρ* (S4 Table).

**Fig 2.**
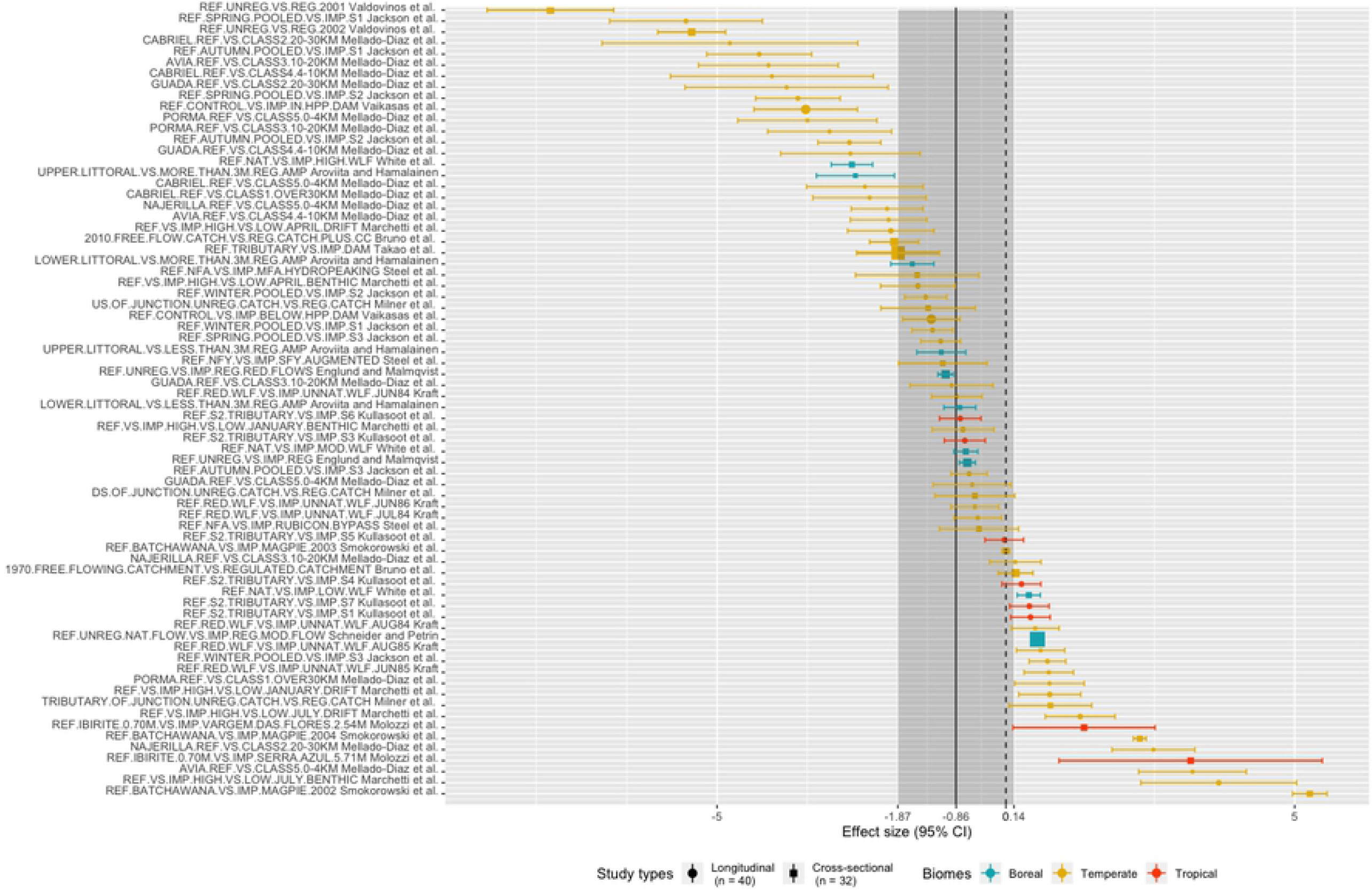
Forest plot of the meta-analysis, the mean effect size is −0.864 (95% CI = −1.87 to 0.144, shaded grey area), where study type is shape-coded (*i*.*e*., circle for longitudinal studies and squares for cross-sectional studies) and biome color coded (*i*.*e*., boreal in blue, temperate in yellow and tropical in red). A negative effect size means that there is a negative impact of hydropower in impacted sites as opposed to reference sites, whereas a positive effect size means that there is positive impact of hydropower in impacted sites as opposed to reference sites.

Moderators had very little influence in the effects of dams and reservoirs on macroinvertebrate richness. Biome did not significantly explain variability effects sizes. For this moderator, it was only possible to make statistically valid inferences for the temperate level (estimate = −0.56; 95% CI = −2.60 to 1.49; p = 0.522; Fig 3a). The boreal and tropical levels had too little degrees of freedom (df_s_ < 4), which invalidates the Satterthwaithe approximation (*i*.*e*., calculation of the effective df_s_ of a linear combination of independent sample variances; [51,53,54]), thus the results of the RVE with this moderator in the equations have to be interpreted with caution. On the contrary, both type of impact and study moderators had enough df_s_ for robust statistical analysis. Statistically significant effects were neither found for downstream flow regulation (estimate = −0.63; 95% CI = −1.53 to 0.28; p = 0.158) nor for upstream water level fluctuations/drawdown impact types (estimate = −0.83; 95% CI = −4.16 to 2.49; p = 0.575; Fig 3b). When study type was used as a moderator, there was no significant difference in effect sizes for cross-sectional design studies (*i*.*e*., natural versus impacted; estimate = 0.49; 95% CI = −1.49 to 2.47; p = 0.581), but there was a marginally significant difference for longitudinal gradient type studies (*i*.*e*., spatial gradient; estimate = −1.21; 95% CI = −2.76 to 0.33; p = 0.095; Fig 3c). Studies that were considered as gradients were most often associated with negative effect sizes. As for results from the season (Fig 3d) and sampling gear (Fig 3e) moderators, no strong statistical inferences can be drawn due to insufficient df_s_ (Satterthwaithe approximation invalidated).

**Fig 3.**
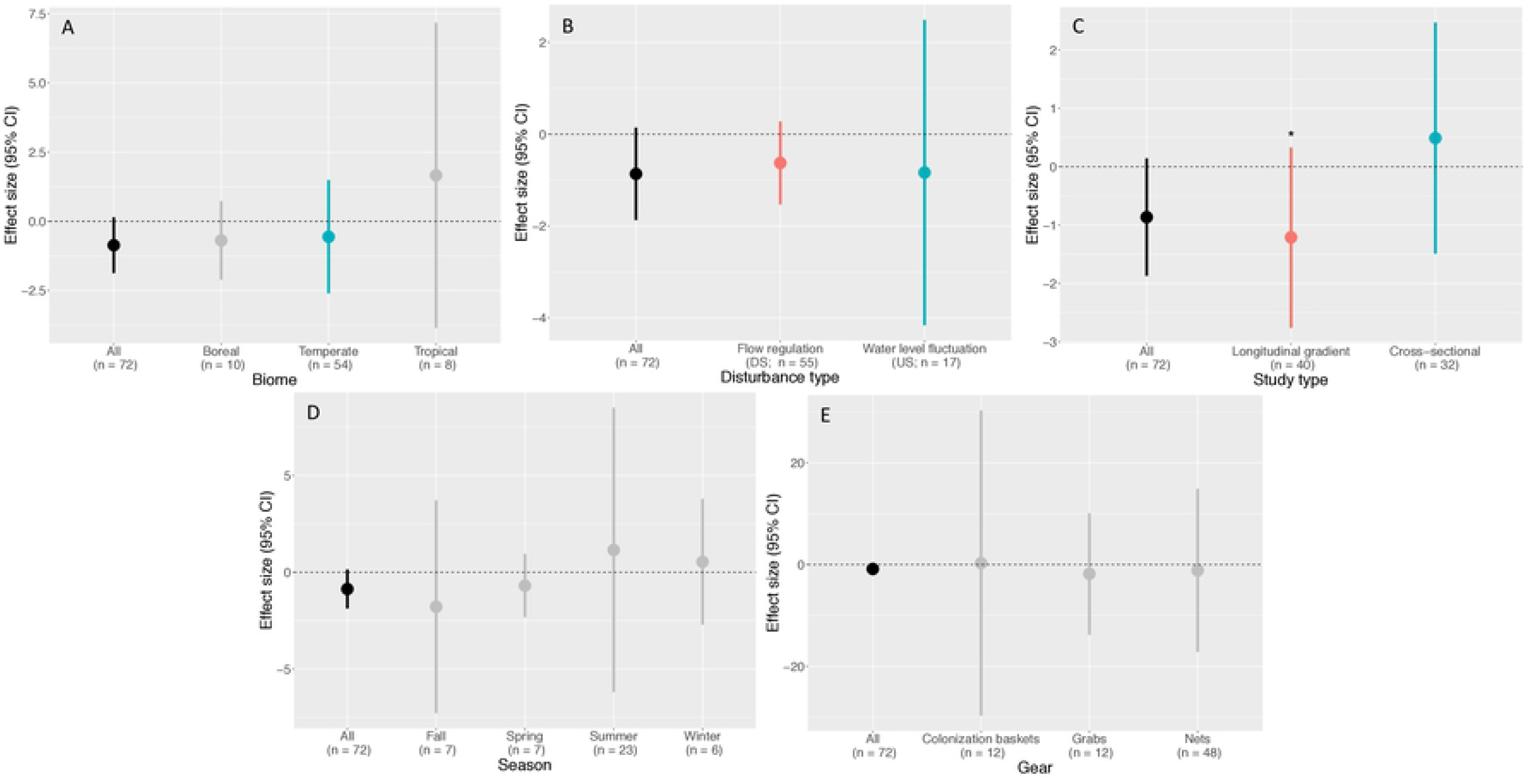
Plots showing the mean effect sizes and their confidence interval for each of the moderators. Value in black is the mean effect size of the meta-analysis and the other colors are related to the different effect sizes when including specific moderators. When in grey, statistical significance of moderator cannot be interpreted with confidence due to statistical power issues (df_s_ insufficient). When effect size is in color (*i*.*e*., blue or red) statistical interpretation can be made with confidence, whether it is significant or not (sufficient df_s_). Asterisk signifies statistically marginally significant effect.

## Discussion

The meta-analysis conducted on 72 pairs of observations (*i*.*e*., impacted versus reference) extracted from 17 studies suggests that hydropower does not statistically impact macroinvertebrate richness in a clear directional way. As highlighted by the literature, there is a divergence in results and it is not obvious as to whether or not hydropower dam, reservoir creation and altered hydrological regime impact macroinvertebrates richness. The lack of statistical significance in the mean effect size seems to result from a large variability in responses, from very negative to very positive of impacts, and not of a real absence of impacts on macroinvertebrate richness (Fig 2). In fact, more than 86% of the effect sizes were significantly different from zero, from which 60% of the observations showed reduced richness due to hydropower and only 26% of the observations showed reduced richness in reference conditions. These percentages, along with the marginally non-significant effect of the mean effect size (*i*.*e*., −0.864; 95% CI = −1.87 to 0.144; p = 0.088) could suggest a tendency toward negative impacts of hydropower on macroinvertebrate richness, this is, however, not supported by statistical evidences.

We observed a large heterogeneity in macroinvertebrate richness responses to hydropower (Fig 2). Part of this variability across observations and studies can usually be explained by environmental and methodological variability, which we can try to control using moderators. In our meta-analysis, we considered biome (*i*.*e*., boreal, temperate or tropical), type of impact (*i*.*e*., water level fluctuations in the reservoir or flow regulation downstream of the dam), the type of study (*i*.*e*., spatial longitudinal gradient [upstream vs downstream] or cross-sectional [natural lake versus impacted reservoir), sampling season (*i*.*e*., spring, summer, fall and winter) and sampling gear (*i*.*e*., net, grab and colonization basket) as moderators. Despite the documented effects on macroinvertebrate richness of the moderators included in our analysis, none of our moderators statistically explained heterogeneity in macroinvertebrate richness responses to hydropower.

### Biome moderator

In general, it is quite common to observe a latitudinal gradient in species richness, where there is a maximum richness in the tropics and a decline towards the poles [31,32,55,56]. In a meta-analysis, Turgeon et al. [57], showed that the impacts of impoundments on fish richness was different across biomes. Significant declines in richness were observed in the tropics, a lower decline was observed in temperate regions and no impacts was observed in boreal biomes. Thus, we hypothesized that a latitudinal gradient could also influence the mean effect size in macroinvertebrates. Our data did not support a latitudinal trend which is not too surprising because there is no clear pattern about whether or not macroinvertebrate richness follows a latitudinal gradient [58–60]. Pearson and Boyero [61] observed a richness peak around the equator for dragonflies (*i*.*e*., odonata), but no clear global pattern for caddisflies (*i*.*e*., trichoptera), whereas Vinson and Hawkins [59] observed richness peaks at mid-latitudes in South and North America for mayflies (*i*.*e*., ephemeroptera), stoneflies (*i*.*e*., plecoptera) and caddisflies (*i*.*e*., trichoptera; [EPT] orders). On the contrary, Scott et al. [62] did not observe this latitudinal gradient for EPT in northern Canada. Geographic range (*i*.*e*., narrow range and exclusion of extreme latitudes; [59]), sampling effort [63], macroinvertebrate specific life-history strategies [61,62] and data resolution (*i*.*e*., weak gradient for local species richness; [64]) were probably the reason behind this lack of consensus [65], and absence of a trend in this meta-analysis. Moreover, the unbalanced sample size across biomes may have impeded any clear conclusions. Boreal samples represented 14% of the data, tropical samples 11% and roughly 75% of the data belong to temperate ecosystems. This prevented us from statistically supporting a trend for both boreal and tropical observations.

### Impact type moderator

Water level fluctuation in reservoirs is characterized by yearly drawdown (*i*.*e*., long-term oscillations in the water level), whereas flow regulation usually is reflected by daily or weekly changes in flow downstream of a dam (*i*.*e*., short-term oscillations; [21]), keeping in mind that variation in the reservoir are also reflected downstream. In reservoir experiencing water level fluctuations in the form of winter drawdown (24% of the reservoirs in this meta-analysis), the littoral zone is exposed to desiccation and freezing for an extended period of time, which has been causing a loss of macroinvertebrate taxa [21,22,66,67] and decreased their overall abundance [68]. In the case of daily downstream fluctuations (76% of the reservoirs in this meta-analysis), organisms can burrow in the sediment (*i*.*e*., low mobility taxa such as oligochaetes) or follow the water level up and down (*i*.*e*., high mobility swimming taxa such as dragonfly nymphs; [21]). Riverine organisms have evolved and developed adaptations to survive flood and drought hydrological dynamics [69], but lentic taxa are not adapted to extreme water level fluctuations, such as winter drawdown in reservoirs (> 2m amplitude; [23]). Because of that, we were expecting a higher impact and effect size of water level fluctuation in reservoir (*i*.*e*., winter drawdown) on macroinvertebrates richness than downstream flow regulation but found no difference nor significant effect of impact type moderator.

### Study type moderator

Another moderator that could explain heterogeneity in our results was the type of study. An ideal way to analyze the impact of hydropower on richness is to set the reference conditions as richness before impoundment versus impacted conditions as richness after impoundment, within the same ecosystem (*e*.*g*., a reference river that was impounded into an impacted reservoir; longitudinal in time, also known as BACI; [33]). However, studies using this type of methodology are rare, even more so for macroinvertebrates, and such studies were mostly absent in the results from the literature review and if present, did not fulfill the required criteria to be included in the meta-analysis (none could be included in this case). Here, we accounted for two types of study methodologies; a cross-sectional methodology (*i*.*e*., comparing a hydropower impacted ecosystem with a natural/reference ecosystem; *n* = 44%) and longitudinal gradient in space methodology (*i*.*e*., upstream of a dam versus downstream of that same dam, within a single ecosystem; *n* = 56%), which are both considered as simple CI studies [33]. These methodologies are not ideal, compared to the longitudinal in time methodology (*i*.*e*., BACI or BA studies), as their reference spatial point differs from the impacted spatial point, thus introducing some environmental noise and there is no way to control for environmental stochasticity [33]. In the cross-sectional type, studies included inter-ecosystem’s variation (*i*.*e*., impacted ecosystem was spatially independent from reference/non-impacted ecosystem). This inherently added some heterogeneity in the responses and the impacts could be more difficult to detect. In the longitudinal in space studies (reference upstream and impacted downstream of the dam, at a single point in time), the observations were from the same ecosystem. Here, there is less heterogeneity and thus, we could have expected the results to be less variable. However, there was no significant difference between the two study types in this meta-analysis. Even though the patterns were not significant, we observed a negative trend in the spatially longitudinal gradient studies (*i*.*e*., higher richness in reference ecosystems, that is upstream of the dam), with a slightly tighter range of variation. The cross-sectional studies had a tendency toward higher richness in reservoirs, with a larger range of variation. This might highlight a problem with general study design and the choice of the reference ecosystem, and caution is in order when interpreting these trends. Nevertheless, we believe that cross-sectional and longitudinal in space references are the best benchmark available to overcome current limitations regarding the lack of richness data before impoundment in the literature.

The analysis of moderators did not allow a better understanding of the large heterogeneity in the effect sizes and suggests that maybe other moderators, which were not available for the studies included in this meta-analysis, could help tease out some of that heterogeneity. For instance, given that macroinvertebrate are very adapted to their localized environmental conditions (*e*.*g*., temperature, granulometry, wave disturbance and macrophytes; [35]), their richness maybe regulated at a much finer scale. Thus, these micro-habitat moderators could be especially relevant to include in a future meta-analysis, although very hard to collect in such a global consolidating endeavour.

## Conclusion

Overall, our meta-analysis did not draw any clear, directional, statistically significant conclusions regarding whether or not hydropower impacts macroinvertebrate richness. However, the large variability observed across studies (significantly negative to significantly positive results), coupled with the marginality of the statistical significance, suggest that macroinvertebrates could be impacted by hydropower, keeping in mind that this suggestion is not statistically supported. The environmental and methodological heterogeneity in the studies might have hindered the detection of a significant effect but none of our moderators helped untangle that heterogeneity. This advises that other moderators, which have not been included in this study due to unavailability among the studies, may be responsible for some of that heterogeneity. We advocate that local, smaller scale variables pertaining to habitat physicochemical characteristics may bring some clarity about the large heterogeneity in effect sizes. As new studies evaluating the impacts of hydropower on macroinvertebrate richness accumulate, we would advise that information regarding local habitat variables be available so they could be recorded and evaluated as moderators in future meta-analyses. This meta-analysis wasn’t able to highlight a clear directional effect of hydropower on macroinvertebrate richness and it would be even more interesting to keep enriching it, as new studies are available, to see if the results could change in a few years and become more assertive. If so, we could finally come to terms with the divergence of results regarding the impacts of hydropower on macroinvertebrate richness and draw clear, statistically supported conclusions.

## Acknowledgements

We thank the CIRAIG – Polytechnique Montréal for covering the publication fees. We also thank the CSBQ for offering systematic reviews and meta-analyses workshops, which were more than useful for putting together this meta-analysis research article.

## Supporting information

**S1 Fig**. PRISMA flow diagram for this meta-analysis from Moher et al. [38].

**S1 Table**. PRISMA checklist for this meta-analysis, from Moher et al. [38].

**S2 Fig**. Funnel plot for this meta-analysis, where no statistically significant asymmetry is observed (z = −1.0707, p = 0.2843).

**S2 Table**. Metadata table showing all variables for each studies included in this meta-analysis *[14,15,17–19,21– 23,26,27,44,70–75]*. Use S3 Table as a companion table to get more information on each variable.

**S3 Table**. Companion table describing all variables in S2 Table.

**S4 Table**. Table showing the sensitive analysis outputs. Rho values (*ρ*) ranges from 0 to 1 and mean effect size (ES), standard error (SE) and between study variance (*τ*^2^) estimates are relatively insensitive to these varying *ρ* values.

## References

[1] Strayer DL, Dudgeon D. Freshwater biodiversity conservation: recent progress and future challenges. J North Am Benthol Soc 2010;29:344–58. https://doi.org/10.1899/08-171.1.

[2] Dudgeon D, Arthington AH, Gessner MO, Kawabata Z-I, Knowler DJ, Leveque C, et al. Freshwater biodiversity: impotance, threats, status and conservation challenges. Biol Rev 2006;81:163–82. https://doi.org/10.1017/S1464793105006950.

[3] Vörösmarty CJ, McIntyre PB, Gessner MO, Dudgeon D, Prusevich A, Green P, et al. Global threats to human water security and river biodiversity. Nature 2010;467:555–61. https://doi.org/10.1038/nature09440.

[4] Gracey EO, Verones F. Impacts from hydropower production on biodiversity in an LCA framework—review and recommendations. Int J Life Cycle Assess 2016;21:412–28. https://doi.org/10.1007/s11367-016-1039-3.

[5] Ward JV, Stanford JA. Ecological connectivity in alluvial river ecosystems and its disruption by flow regulation. Regul Rivers Res Manag 1995;11:105–19. https://doi.org/10.1002/rrr.3450110109.

[6] Kummu M, Varis O. Sediment-related impacts due to upstream reservoir trapping, the Lower Mekong River. Geomorphology 2007;85:275–93. https://doi.org/10.1016/j.geomorph.2006.03.024.

[7] Nilsson C, Brown RL, Jansson R, Merritt DM. The role of hydrochory in structuring riparian and wetland vegetation. Biol Rev 2010;85:837–58.

[8] Poff NL, Allan JD, Bain MB, Karr JR, Prestegaard KL, Richter BD, et al. The Natural Flow Regime. BioScience 1997;47:769–84. https://doi.org/10.2307/1313099.

[9] Bunn SE, Arthington AH. Basic Principles and Ecological Consequences of Altered Flow Regimes for Aquatic Biodiversity. Environ Manage 2002;30:492–507. https://doi.org/10.1007/s00267-002-2737-0.

[10] Friedl G, Wüest A. Disrupting biogeochemical cycles - Consequences of damming. Aquat Sci 2002;64:55– 65. https://doi.org/10.1007/s00027-002-8054-0.

[11] Santucci VJ, Gephard SR, Pescitelli SM. Effects of Multiple Low-Head Dams on Fish, Macroinvertebrates, Habitat, and Water Quality in the Fox River, Illinois. North Am J Fish Manag 2005;25:975–92. https://doi.org/10.1577/M03-216.1.

[12] Poff NL, Olden JD, Merritt DM, Pepin DM. Homogenization of regional river dynamics by dams and global biodiversity implications. Proc Natl Acad Sci 2007;104:5732–7. https://doi.org/10.1073/pnas.0609812104.

[13] Liermann CR, Nilsson C, Robertson J, Ng RY. Implications of Dam Obstruction for Global Freshwater Fish Diversity. BioScience 2012;62:539–48. https://doi.org/10.1525/bio.2012.62.6.5.

[14] Jackson HM, Gibbins CN, Soulsby C. Role of discharge and temperature variation in determining invertebrate community structure in a regulated river. River Res Appl 2007;23:651–69. https://doi.org/10.1002/rra.1006.

[15] Takao A, Kawaguchi Y, Minagawa T, Kayaba Y, Morimoto Y. The relationships between benthic macroinvertebrates and biotic and abiotic environmental characteristics downstream of the Yahagi Dam, Central Japan, and the State Change Caused by inflow from a Tributary. River Res Appl 2008;24:580–97. https://doi.org/10.1002/rra.1135.

[16] Behrend RDL, Takeda AM, Gomes LC, Fernandes SEP. Using oligochaeta assemblages as an indicator of environmental changes. Braz J Biol 2012;72:873–84. https://doi.org/10.1590/S1519-69842012000500014.

[17] Kullasoot S, Intrarasattayapong P, Phalaraksh C. Use of benthic macroinvertebrates as bioindicators of anthropogenic impacts on water quality of Mae Klong river, Western Thailand 2017.

[18] Kraft KJ. Effect of Increased Winter Drawdown on Benthic Macroinvertebrates in Namakan Reservoir, Voyeurs National Park. Houghton, Michigan: Michigan Techonological University; 1988.

[19] Englund G, Malmqvist B. Effects of Flow Regulation, Habitat Area and Isolation on the Macroinvertebrate Fauna of Rapids in North Swedish Rivers. Regul Rivers Res Manag 1996;12:433–45. https://doi.org/10.1002/(SICI)1099-1646(199607)12:4/5<433::AID-RRR415>3.0.CO;2-6.

[20] Malmqvist B, Englund G. Effects of hydropower-induced flow perturbations on mayfly (Ephemeroptera) richness and abundance in north Swedish river rapids. Hydrobiologia 1996;341:145–58. https://doi.org/10.1007/BF00018118.

[21] Valdovinos C, Moya C, Olmos V, Parra O, Karrasch B, Buettner O. The importance of water-level fluctuation for the conservation of shallow water benthic macroinvertebrates: an example in the Andean zone of Chile. Biodivers Conserv 2007;16:3095–109. https://doi.org/10.1007/s10531-007-9165-7.

[22] Aroviita J, Hämäläinen H. The impact of water-level regulation on littoral macroinvertebrate assemblages in boreal lakes. Hydrobiologia 2008;613:45–56. https://doi.org/10.1007/s10750-008-9471-4.

[23] White MS, Xenopoulos MA, Metcalfe RA, Somers KM. Water level thresholds of benthic macroinvertebrate richness, structure, and function of boreal lake stony littoral habitats. Can J Fish Aquat Sci 2011;68:1695–704. https://doi.org/10.1139/f2011-094.

[24] Głowacki Ł, Grzybkowska M, Dukowska M, Penczak T. Effects of damming a large lowland river on chironomids and fish assessed with the (multiplicative partitioning of) true/Hill biodiversity measure. River Res Appl 2011;27:612–29. https://doi.org/10.1002/rra.1380.

[25] Serafini Floss EC, Secretti E, Kotzian CB, Spies MR, Pires MM. Spatial and temporal distribution of non-biting midge larvae assemblages in streams in a mountainous region in southern Brazil. J INSECT Sci 2013;13.

[26] Smokorowski KE, Metcalfe RA, Finucan SD, Jones N, Marty J, Power M, et al. Ecosystem level assessment of environmentally based flow restrictions for maintaining ecosystem integrity: a comparison of a modified peaking versus unaltered river. Ecohydrology 2011;4:791–806. https://doi.org/10.1002/eco.167.

[27] Marchetti MP, Esteban E, Smith ANH, Pickard D, Richards AB, Slusark J. Measuring the ecological impact of long-term flow disturbance on the macroinvertebrate community in a large Mediterranean climate river. J Freshw Ecol 2011;26:459–80. https://doi.org/10.1080/02705060.2011.577974.

[28] De Palma A, Sanchez-Ortiz K, Martin PA, Chadwick A, Gilbert G, Bates AE, et al. Chapter Four - Challenges With Inferring How Land-Use Affects Terrestrial Biodiversity: Study Design, Time, Space and Synthesis. In: Bohan DA, Dumbrell AJ, Woodward G, Jackson M, editors. Adv. Ecol. Res., vol. 58, Academic Press; 2018, p. 163–99. https://doi.org/10.1016/bs.aecr.2017.12.004.

[29] Viechtbauer W. Conducting Meta-Analyses in with the Package. J Stat Softw 2010;36. https://doi.org/10.18637/jss.v036.i03.

[30] Gibson L, Lee TM, Koh LP, Brook BW, Gardner TA, Barlow J, et al. Primary forests are irreplaceable for sustaining tropical biodiversity. Nature 2011;478:378–81. https://doi.org/10.1038/nature10425.

[31] Rosenzweig ML. Species diversity in space and time. Cambridge, U.K.: Cambridge University Press; 1995.

[32] Willig MR, Kaufman DM, Stevens RD. Latitudinal Gradients of Biodiversity: Pattern, Process, Scale, and Synthesis. Annu Rev Ecol Evol Syst 2003;34:273–309. https://doi.org/10.1146/annurev.ecolsys.34.012103.144032.

[33] Christie AP, Amano T, Martin PA, Shackelford GE, Simmons BI, Sutherland WJ. Simple study designs in ecology produce inaccurate estimates of biodiversity responses. J Appl Ecol 2019;56:2742–54. https://doi.org/10.1111/1365-2664.13499.

[34] Hill MJ, Sayer CD, Wood PJ. When is the best time to sample aquatic macroinvertebrates in ponds for biodiversity assessment? Environ Monit Assess 2016;188. https://doi.org/10.1007/s10661-016-5178-6.

[35] McCafferty WP. Aquatic entomology: The fishermen’s and ecologists’ illustrated guide to insects and their relatives. Jones & Bartlett Learning; 1983.

[36] Wetzel RG. Limnology: Lake and river ecosystems. California: Gulf Professional Publishing; 2001.

[37] Koricheva J, Gurevitch J, Mengersen K. Handbook of Meta-Analysis in Ecology and Evolution. Princeton and Oxford: Princeton University Press; 2013.

[38] Moher D. Preferred Reporting Items for Systematic Reviews and Meta-Analyses: The PRISMA Statement. Ann Intern Med 2009;151:264. https://doi.org/10.7326/0003-4819-151-4-200908180-00135.

[39] Clarivate Analytics. Browse, search, and explore journals indexed in the Web of Science. Web Sci Group Clarivate Anal Co 2019. https://mjl.clarivate.com/ (accessed November 27, 2019).

[40] Clarivate Analytics. Web of Science Core Collection Help 2018. http://images.webofknowledge.com/WOKRS533JR18/help/WOS/hs_sort_options.html (accessed November 27, 2019).

[41] Olson DM, Dinerstein E, Wikramanayake ED, Burgess ND, Powell GVN, Underwood EC, et al. Terrestrial Ecoregions of the World: A New Map of Life on Earth. BioScience 2001;51:933–8.

[42] Mitchell M, Muftakhidinov B, Winchen T. Engauge Digitizer Software. 2017.

[43] Borenstein M, Hedges LV, Higgins JPT, Rothstein HR. Introduction to Meta-Analysis. John Wiley & Sons; 2011.

[44] Schneider SC, Petrin Z. Effects of flow regime on benthic algae and macroinvertebrates - A comparison between regulated and unregulated rivers. Sci Total Environ 2017;579:1059–72. https://doi.org/10.1016/j.scitotenv.2016.11.060.

[45] Hedges LV. Distribution Theory for Glass’s Estimator of Effect size and Related Estimators, Distribution Theory for Glass’s Estimator of Effect size and Related Estimators. J Educ Stat 1981;6:107–28. https://doi.org/10.3102/10769986006002107.

[46] Wickham H. ggplot2: Elegant Graphics for Data Analysis. New York: Springer-Verlag; 2016.

[47] Sterne JAC, Egger M. Funnel plots for detecting bias in meta-analysis: Guidelines on choice of axis. J Clin Epidemiol 2001;54:1046–55. https://doi.org/10.1016/S0895-4356(01)00377-8.

[48] Sterne JAC, Egger M. Regression methods to detect publication and other bias in meta-analysis. In: Rothstein HR, Sutton AJ, Borenstein M, editors. Publ. Bias Meta-Anal. Prev. Assess. Adjust. 1 st ed., Chichester (UK): Wiley; 2005, p. 99–110.

[49] Huedo-Medina TB, Sánchez-Meca J, Marín-Martínez F, Botella J. Assessing heterogeneity in meta-analysis: Q statistic or I2 index? Psychol Methods 2006;11:193–206. https://doi.org/10.1037/1082-989X.11.2.193.

[50] Moeyaert M, Ugille M, Beretvas SN, Ferron J, Bunuan R, Noortgate WV den. Methods for dealing with multiple outcomes in meta-analysis: a comparison between averaging effect sizes, robust variance estimation and multilevel meta-analysis. Int J Soc Res Methodol 2017;20:559–72. https://doi.org/10.1080/13645579.2016.1252189.

[51] Fisher Z, Tipton E. robumeta: an R package for robust variance estimation in meta-analysis 2017:16.

[52] R Core Team. R: a language and environment for statistical computing. Vienna, Austria: R Foundation for Statistical Computing; 2017.

[53] Emden H van. Statistics for Terrified Biologists. Wiley; 2009.

[54] Spellman FR, Whiting NE. Handbook of Mathematics and Statistics for the Environment. CRC Press; 2013.

[55] Hillebrand H. On the Generality of the Latitudinal Diversity Gradient. Am Nat 2004;163:192–211. https://doi.org/10.1086/381004.

[56] Lomolino MV, Riddle BR, Brown JH. Biogeography. 3rd ed. Sunderland, MA: Sinauer Associates; 2006.

[57] Turgeon K, Turpin C, Gregory-Eaves I. Dams have varying impacts on fish communities across latitudes: a quantitative synthesis. Ecol Lett 2019;22:1501–16. https://doi.org/10.1111/ele.13283.

[58] Covich AP. Geographical and Historical Comparisons of Neotropical Streams: Biotic Diversity and Detrital Processing in Highly Variable Habitats. J North Am Benthol Soc 1988;7:361–86. https://doi.org/10.2307/1467297.

[59] Vinson MR, Hawkins CP. Broad-scale geographical patterns in local stream insect genera richness. Ecography 2003;26:751–67. https://doi.org/10.1111/j.0906-7590.2003.03397.x.

[60] Allan JD, Castillo MM. Stream Ecology: Structure and function of running waters. Springer Science & Business Media; 2007.

[61] Pearson RG, Boyero L. Gradients in regional diversity of freshwater taxa. J North Am Benthol Soc 2009;28:504–14. https://doi.org/10.1899/08-118.1.

[62] Scott RW, Barton DR, Evans MS, Keating JJ. Latitudinal gradients and local control of aquatic insect richness in a large river system in northern Canada. J North Am Benthol Soc 2011;30:621–34. https://doi.org/10.1899/10-112.1.

[63] Vinson MR, Hawkins CP. Effects of Sampling Area and Subsampling Procedure on Comparisons of Taxa Richness among Streams. J North Am Benthol Soc 1996;15:392–9. https://doi.org/10.2307/1467286.

[64] Heino J. Biodiversity of Aquatic Insects: Spatial Gradients and Environmental Correlates of Assemblage-Level Measures at Large Scales. Freshw Rev 2009;2:1–29. https://doi.org/10.1608/FRJ-2.1.1.

[65] Shah DN, Domisch S, Pauls SU, Haase P, Jähnig SC. Current and future latitudinal gradients in stream macroinvertebrate richness across North America. Freshw Sci 2014;33:1136–47. https://doi.org/10.1086/678492.

[66] Palomäki R, Koskenniemi E. Effects of bottom freezing on macrozoobenthos in the regulated lake Pyhäjärvi. Arch Für Hydrobiol 1993;128:73–90.

[67] Palomäki R. Response by macrozoobenthos biomass to water level regulation in some Finnish lake littoral zones. Hydrobiologia 1994;286:17–26. https://doi.org/10.1007/BF00007277.

[68] Trottier G, Embke H, Turgeon K, Solomon C, Nozais C, Gregory-Eaves I. Macroinvertebrate abundance is lower in temperate reservoirs with higher winter drawdown. Hydrobiologia 2019;834:199–211. https://doi.org/10.1007/s10750-019-3922-y.

[69] Humphries P, Baldwin DS. Drought and aquatic ecosystems: an introduction. Freshw Biol 2003;48:1141–6. https://doi.org/10.1046/j.1365-2427.2003.01092.x.

